# Histone H3K4me3 and H3K9me3 are super over-methylated in soft tissue sarcoma compared to normal muscle in patient-derived xenograft (PDX) mouse models: an indicator of cancer methionine addiction

**DOI:** 10.1101/2021.01.01.425040

**Authors:** Yusuke Aoki, Jun Yamamoto, Yasunori Tome, Kazuyuki Hamada, Sachiko Inubushi, Yoshihiko Tashiro, Michael Bouvet, Itaru Endo, Kotaro Nishida, Robert M. Hoffman

## Abstract

Methionine addiction is a fundamental and general hallmark of cancer discovered by us almost a half-century ago [Proc Natl Acad Sci U S A 73 (1976) 1523-1527]. Methionine addiction is defined as the requirement, specific for cancer cells of all types, for exogenous methionine despite the normal ability to synthesize methionine from homocysteine. The methionine addiction of cancer is termed the Hoffman-effect, analogous to the Warburg-effect of the high glucose requirement of cancer cells. Methionine addiction is due to excess transmethylation reactions resulting in high methionine flux in cancer cells, which causes them to selectively arrest under methionine restriction due to depletion of free methionine and S-adenosyl methionine. Recently we have shown methionine-addicted cancer cells over-methylate histone H3 lysine marks which are not over-methylated in normal cells or in low-malignancy methionine-independent revertants derived from methionine-addicted cancer cells. In the present report, we show that in patient-derived xenograft (PDX) mouse models of the most common soft tissue sarcomas: myxofibrosarcoma, undifferentiated pleomorphic sarcoma (UPS) and liposarcoma, histone H3K4me3 and H3K9me3 are super over-methylated compared to normal muscle tissue. This new result is discussed along with our previous reports, regarding the potential of histone H3 over-methylation as a basis of malignancy.

## 1. Introduction

The paradigm for the basis of cancer has changed drastically over the past century. In the early 20th century, aneuploidy was proposed as the basis of cancer [1] and is still thought so by some researchers [2]. In the 1920’s a German-Jewish biochemist Otto Warburg proposed that the high rate of aerobic glycolysis, due to defective mitochondria, was the basis of cancer. This was termed the Warburg-effect [3]. In the 1960’s another paradigm change took place, stating that viruses are the cause of cancer [4]. After about 15years, the viral-cause-of-cancer paradigm failed, due to inability to find viruses in patient cancers, despite billions of dollars spent. Then in the late 1970’s a new field of cellular oncogenes appeared, with the discovery that certain cancer cells contained mutated genes that could be transferred by transfection to induce greater malignancy in recipient cells [5,6]. The cellular oncogenes were analogous to genes of viruses that could cause cancer in animals, the most famous of which is the Rous sarcoma virus which contains the “Src” gene, discovered by Duesberg and Vogt [7]. The idea of anti-oncogenes or tumor suppressor genes was part of this paradigm [8]. It was thought that when an oncogene or tumor suppressor gene was mutated in a normal cell, the cell became cancerous. However, it became known through more and more sophisticated DNA sequencing technology that cancers had 100s if not 1000s of mutated genes. Presently, this is explained that despite the enormous numbers of mutated and misregulated genes in a particular cancer, the cancer contains a mutated “driver” gene among the set of altered gene that makes it malignant [9]. However, no specific driver gene is consistently found in the various cancers.

However, there is another more general explanation for the basis of cancer, the cancer-specific property of methionine addiction, discovered by one of us (RMH), almost a half century ago [10]. Methionine addiction is the requirement of cancer cells for exogenous methionine, despite their ability to make normal or super-normal amounts of methionine from homocysteine. In contrast normal cells or methionine-independent revertants derived from methionine-addicted cancer cells require much less methionine [10–14]. It was then found in the 1980s that methionine addiction was due to high flux of methionine in cancer cells, due to excess transmethylation reactions [15] which deplete free methionine and S-adenosyl methionine in cancer cells during methionine restriction (MR) but not in normal cells or methionine-independent revertants [10,12–14]. Methionine addiction is termed the Hoffman effect [16] analogous to the Warburg effect of glucose addiction of cancer [3]. However, for 35 years the fate of transferred methyl groups in cancer cells was unknown. In 2019, a major hint toward solving the problem of methionine addiction came from the observation that “tumor-initiating cells” have highly methylated histone H3 lysine marks [17]. We then observed that cancer cell lines have highly methylated histone H3 lysine marks that are absent in normal cells and in low-malignancy methionine independence revertant derived from the methionine-addicted cancer cells [18]. In addition we have found that over-methylated histone H3 lysine marks are unstable in methionine addicted cancer cells under methionine restriction [19] in which condition they arrest in late-S / G_2_ of the cell cycle [20,21]. In contrast, normal cells have a normal level of methylated histone H3 lysine marks which are stable under methionine restriction [19].

In the present report, we show that the most common types of soft tissue sarcoma (STS), myxofibrosarcoma, undifferentiated pleomorphic sarcoma (UPS) and liposarcoma, growing in patient derived xenograft (PDX) mouse models, have super over-methylated histone H3K4me3 and H3K9me3 marks, in contrast to normal muscle in the host nude mouse. The results of the present report, along with our previous results [18,19], are discussed as the possible basis of malignancy.

## 2. Materials and Methods

### 2.1. Mice

Athymic nu/nu nude mice (AntiCancer, Inc., San Diego, CA, USA) were used to establish PDX mouse models of three types of STS: myxofibrosarcoma, undifferentiated pleomorphic sarcoma (UPS), liposarcoma. All studies were performed with a protocol approved by an AntiCancer, Inc. Institutional Animal Care and Use Committee (IACUC) following the principles and procedures provided in the National Institutes of Health (NIH) Guide for the Care and Use of Animals under Assurance Number A3873-1. In order to minimize any suffering of the animals, anesthesia and analgesics were used for all surgical experiments.

### 2.2. Patient-derived tumors

#### 2.2-1. Myxofibrosarcoma

A resected tumor tissue specimen of a 64-year-old woman with right anterior thigh myxofibrosarcoma was previously obtained from surgery at the UCLA medical center and transported immediately to the laboratory at AntiCancer, Inc.. This sample was cut into 5 mm^3^ fragments and then implanted subcutaneously in nude mice as previously reported [22]

#### 2.2-2. undifferentiated pleomorphic sarcoma (UPS)

A resected tumor tissue specimen of a 65-year-old man with a right thigh UPS obtained at surgery in UCLA was previously brought to AntiCancer, Inc. and implanted subcutaneously in nude mice as previously described. [23]

#### 2.2-3. Liposarcoma

A tumor specimen resected from a 67-year-old man with upper abdominal softtissue liposarcoma at UCLA, was brought to Anticancer, Inc. and implanted subcutaneously in nude mice as previously described [24].

All three sarcoma cases were approved by the UCLA Institutional Review Board (IRB#10-001857) and written informed consent was obtained from all patients.

### 2.3. Immunoblotting

The sarcoma PDX mouse models were sacrificed and tumor and femoral muscle tissues were resected before tumors reached a volume of 4000 mm^3^. Then tissues were frozen immediately after resections in liquid nitrogen. To prepare for immunoblotting, the tumors were homogenized and the histones were extracted using a Epiquik Total Histone Extraction Kit (Epigentek, Farmingdale, NY, USA). Immunoblotting for these histones was performed under the following conditions: 12% SDS-PAGE gels and 0.2 μm polyvinylidene difluoride (PVDF) membranes were used. Blocking was performed using the Bullet Blocking One for Western Blotting (Nakalai Tesque, Inc. Kyoto, Japan). Anti-H3K4me3 antibody (1:1,000, #9751, Cell Signaling Technology, Danvers, MA, USA); anti-H3K9me3 antibody (1:1,000, #13969, Cell Signaling Technology); and anti-H3 antibody (1:1,500, 17168-1-AP, Proteintech, Rosemont, IL, USA) were used. Total histone H3 was used as a loading control. Horseradish-peroxidase-conjugated anti-rabbit IgG (1:20,000, SA00001-2, Proteintech, Rosemont, IL, USA) was used as a second antibody. Immunoreactivity was visualized using Clarity Western ECL Substrate (Bio-Rad Laboratories, Hercules, CA, USA). The signals were detected with UVP ChemStudio (Analytik Jena, Upland, CA, USA) [18,19].

### 2.4. Hematoxylin and eosin (H & E) staining

After resection of the sarcoma and femoral muscle tissues from PDX mice, the tissues were fixed with 10% formalin overnight before embedding in paraffin. The tumor and muscle tissues were sectioned and specimens were deparaffinized with xylene and rehydrated in a series of ethanol. Staining of H&E was performed in accordance with standard protocol.

### 2.5 immunohistochemical (IHC) staining

The same procedure as for H & E staining, described above, was performed through rehydration. Autoclaving was then performed in order to retrieve antigen at 120 °C for 5 min with a citrate buffer solution (pH 6.0). Next, the specimens were immersed in absolute methanol containing 0.3% hydrogen peroxide solution in order to block endogenous peroxidase activity at room temperature for 30 minutes. Then, the specimens were blocked using 10 % bovine serum albumin at room temperature for 15 min and incubated with H3K4me3 antibody (1:1,000) at 4 °C overnight. IHC reactions were detected with HistoFine (Nichirei, Tokyo, Japan) and Dako Omnis (Agilent, Santa Clara, CA, USA). Finally, the specimens were counterstained with hematoxylin [18,19].

## 3. Results

### 3.1. Histone H3K4me3 and H3K9me3 in the PDX sarcoma tissues are super over-methylated compared to normal muscle

We evaluated the methylation status of histone marks H3K4me3 and H3K9me3 of the myxofibrosarcoma, UPS and liposarcoma grown in their corresponding PDX models. Histone H3K4me3 and H3K9me3 were super over-methylated in all of the three sarcomas compared to muscle tissue from the host mice as seen by immunoblotting (Fig. 1). Super over-methylation of H3K4me3 in all three sarcoma models can also be seen by immunohistochemistry (Fig. 2)

**Figure 1.**
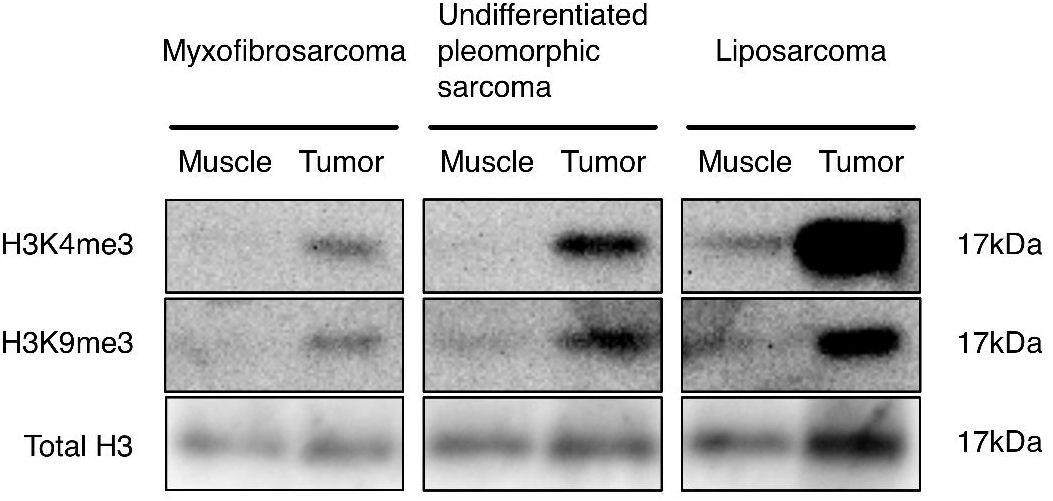
Immunoblotting of histone marks H3K4me3 and H3K9me3 from myxofibrosarcoma, undifferentiated pleomorphic sarcoma and liposarcoma and normal mouse muscle from PDX mouse models. Please see Material and Methods for details.

**Figure 2.**
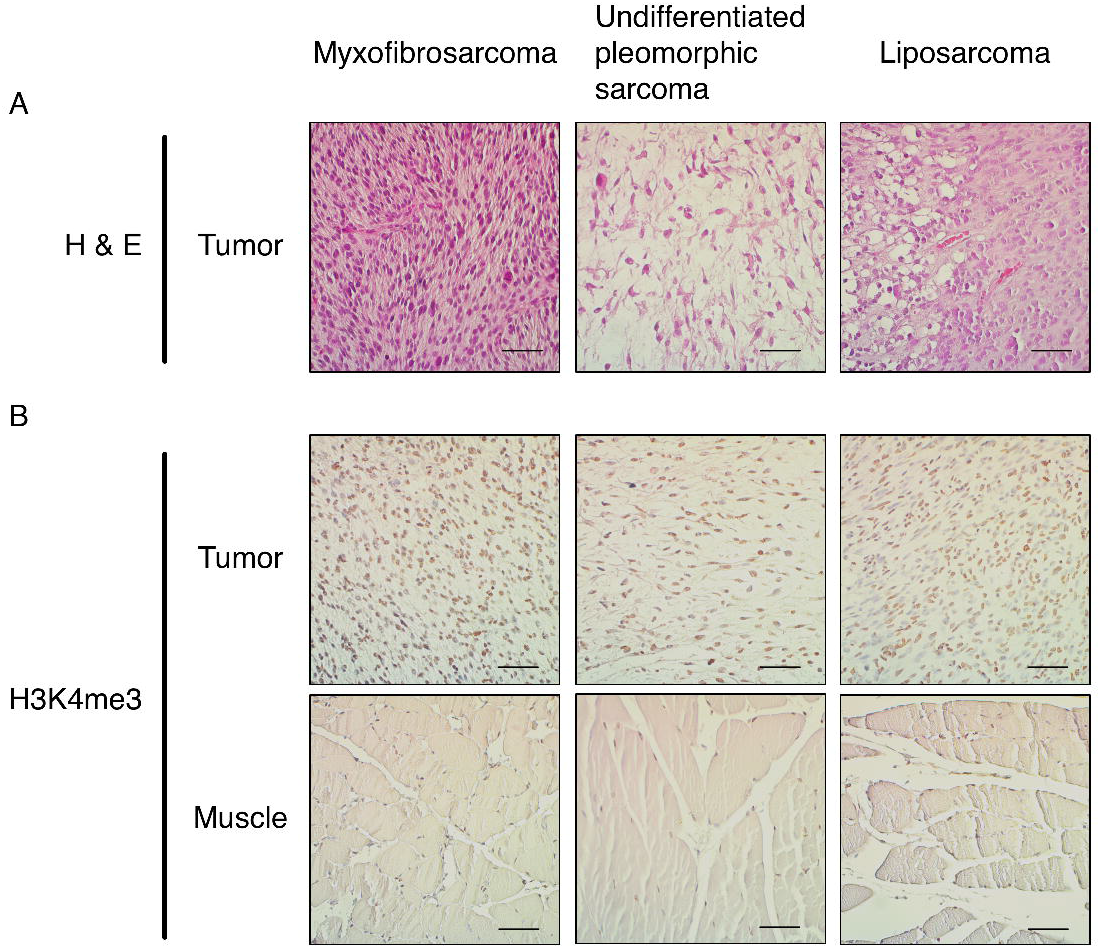
(A) Representative H & E staining images of myxofibrosarcoma, undifferentiated pleomorphic sarcoma and liposarcoma grown in PDX mouse models. (B) Immunohistochemical staining for the histone marks H3K4me3 in a myxofibrosarcoma, undifferentiated pleomorphic sarcoma and liposarcoma grown in PDX mouse models, and normal mouse muscle. Magnification: 200×. Scale bar: 50μm. Please see Material and Methods for details.

### 3.2. Histology of each sarcoma type in the PDX model

H & E staining showed that all of three types of sarcoma have each maintained the histological characteristics of their sarcoma type after growth in PDX mouse models (Fig. 2). The myxofibrosarcoma tissue comprised high-density spindle-shaped cancer cells with myxoid stroma. The UPS tissue comprised high-density pleomorphic cells. The liposarcoma tissue comprised both well differentiated and dedifferentiated cells.

## 4. Discussion

The present study showed that histone marks H3K4me3 and H3K9me3 were super over-methylated in myxofibrosarcoma, UPS and liposarcoma, growing as patient derived xenografts (PDX) nude mice, in contrast to normal muscle from the host mice which was only slightly methylated in histone H3K4me3 and H3K9me3 (Fig. 1). This result is consistent with our previous report that histone H3K4me3 and H3K9me3 are highly methylated in methionine-addicted cancer cell lines but not in normal cells or in low-malignancy methionine-independent revertants derived from the methionine-addicted cell lines [18,19]. This is the first report to our knowledge that histone H3K4me3 and/or H3K9me3 are super over-methylated in patient tumors growing in a PDX model.

Methionine addiction has been found in all properly-tested cancer cell types in vitro [25,26] and vivo [27]. The present results of super over-methylated histone H-3 marks in the three sarcoma PDX types indicate they are also methionine addicted. The present report and our previous reports [18,19] suggest that super over-methylation of histone H3K4me4 and H3K9me3 may be a general biomarker in cancer, since histone H3 lysine over-methylation is linked to methionine addiction, which is found in the all cancer types [25,26].

The enzyme methioninase has been used to effectively target methionine addiction and inhibit or arrest tumor growth in vivo [28–32].

Histone H3K4me3 and H3K9me3 are thought to be global gene regulators and their super over-methylation status may inhibit their function and change the expression of a large number of genes that result in malignant conversion of the cell.

There have been a number of publications in recent years [33–35] attempting to correlate the extent of methylation of histone H3 lysine marks with patient outcome in cancers of the liver, cervical and colon, with conflicting results.

The results of the present study showing super over-methylation of histone H3K4me3 and H3K9me3 in the three types of the most common sarcomas: myxofibrosarcoma, UPS and liposarcoma in PDX mouse models, along with our previous results [18,19], suggest that super over-methylation of histone marks may be an important clinical indicator and that histone H3 super over-methylation may be a basis of malignancy. PET imaging of cancer with [^11^C]methionine gives a very strong signal compared to [^18^F]FDG PET [36,37], indicating that clinical cancer is methionine addicted and that the Hoffman effect is stronger than the Warburg effect.

Future studies will correlate the malignant behavior, including metastasis of the sarcomas, in patient-derived orthotopic xenograft (PDOX) models [38], The present results give additional support that methionine addiction, due at least in part to super over-methylation of H3K4me3 and H3K9me3, is the fundamental hallmark of cancer.

## Author contribution

YA, JY, YT and RMH were involved in study conception and design. YA, JY, KH and RMH were involved in acquisition of data. YA, JY, YT, KH, SI, MB, IE, KN and RMH analyzed and interpreted data. YA, YT and RMH wrote this manuscript. All authors reviewed and approved the manuscript.

